# Human cell line tropism, interferon system interactions and NSs mechanisms by the Ntepes and Gabek Forest phleboviruses

**DOI:** 10.1101/2025.10.17.683117

**Authors:** Matthew J. Pickin, Ulrike Felgenhauer, David P. Tchouassi, Sandra Junglen, Friedemann Weber

**Affiliations:** Institute of Virology, FB10-Veterinary Medicine, Justus-Liebig University, D-35392 Giessen, Germany; International Centre of Insect Physiology and Ecology, Nairobi, Kenya; Institute of Virology, Charité – Universitätsmedizin Berlin, corporate member of Freie Universität Berlin, Humboldt-Universität zu Berlin and Berlin Institute of Health, Berlin, Germany

**Keywords:** Ntepes virus, Gabek Forest virus, human cell line tropism, interferon induction, interferon sensitivity, interferon antagonism, NSs protein, MAVS, CARD

## Abstract

*Phleboviruses* are a large genus within the family *Phenuiviridae,* class *Bunyaviricetes*. These arboviruses are present all over the world, and several members can cause severe disease. New viruses are continuously discovered, but rarely isolated and analysed in cell culture to assess their risk to humans. Here, we describe the *in cellulo* characteristics for two closely related African phleboviruses, Ntepes virus (NTPV) and Gabek Forest virus (GFV), for which human seroprevalence has been reported but pathogenicity remains unknown. For both viruses, human cell lines from liver, lung, and kidney were permissive, and their capacity to suppress and cope with the antiviral type I interferon (IFN) system was comparable to Rift Valley fever phlebovirus MP-12. Consequently, their non-structural protein NSs, a well-known virulence factor of *Phenuiviridae*, was able to interfere with both the induction and signaling of IFN. Blockade of IFN induction is a conserved NSs activity, and we found that in the case of NTPV and GFV, the NSs targets MAVS (Mitochondrial antiviral signaling protein), a cellular signaling adapter which plays a key role in the induction of IFN gene expression. NTPV and GFV NSs thereby strongly and specifically bind the N-terminal CARD (Caspase recruitment domain) of human MAVS, thus occupying the very domain that is required for MAVS to relay the signal coming from the virus sensor RIG-I. Thus, the cell line tropism, IFN system interactions and NSs mechanisms suggest that NTPV and GFV fulfill the *in cellulo* criteria for successful infection of humans.

**Importance:** Chikungunyna virus and Zika virus are just two recent examples of arboviruses that have emerged and caused outbreaks in humans. We investigated whether Ntepes virus (NTPV) and Gabek Forest virus (GFV), two closely related bunyaviruses of the genus *Phlebovirus* with human seroprevalence in certain African regions, have cell culture characteristics that are compatible with a risk for humans. Both viruses were able to infect human cell lines derived from inner organs, and could circumvent the human antiviral interferon system similar to strain MP-12 of the established Rift Valley fever phlebovirus. Specifically, we identified that the interferon antagonists NSs of NTPV and GFV could strongly and specifically bind to a domain in the cellular adaptor protein MAVS that is crucial for the upregulation of the antiviral interferon system. NTPV and GFV may thus have potential to further adapt to humans.

## Introduction

Members of the genus *Phlebovirus* (family *Phenuiviridae,* class *Bunyaviricetes*) are globally abundant arboviruses of which several are known to cause human disease (1) (2–4). Their virions are enveloped and contain the glycoproteins Gn and Gc in the membrane. The viral genome is comprised of three RNA segments (L, M, S) with negative-sense or ambisense polarity that are packaged by the viral nucleocapsid (N) protein and associated with the RNA- dependent RNA polymerase (L). The large (L) segment of the genome codes for the RNA- dependent RNA polymerase (RdRp), the medium (M) segment typically harbours the open reading frame for Gn and Gc, as well as for a medium non-structural protein NSm, and the small (S) segment encodes the nucleoprotein N in negative sense, as well as the non-structural protein NSs in ambisense orientation. NSs is the main virulence factor of phleboviruses, counteracting the antiviral interferon (IFN) system of mammalian hosts (5–7).

Most phleboviruses are transmitted either by mosquitoes or by sandflies (1, 8–10). Rift Valley fever virus (RVFV) is a prominent representative of the mosquito-borne phleboviruses, causing recurrent epidemics in Africa and the Middle East (11–13). In ruminants, the infection leads to abortion storms and often death of young animals; in humans RVFV causes a febrile illness that can progress to hepatitis, retinitis, encephalitis or fatal hemorrhagic fevers, resulting in up to 20% mortality among hospitalized patients (14, 15).

Sandfly fever Sicilian virus (SFSV) and Toscana virus (TOSV) are well-known examples of sandfly-borne phleboviruses that are prevalent in the Mediterranean area and beyond. Human infection can manifest itself as a self-limiting flu-like febrile illness (3-day fever, “dog disease”), and in case of TOSV develop into neuroinvasive disease (1, 8, 10).

Novel phleboviruses are continuously discovered by isolation from vector or host organisms, or by metagenomics analyses (3, 9, 10, 16–20). In rare instances, phleboviruses could be isolated from human cases (21–23). More often, human infection is indicated by seroprevalence studies (24–29), but their implications for actual disease remain unclear. In 2014, a sandfly surveillance study was conducted in Kenya that led to the isolation of a previously unknown phlebovirus termed Ntepes virus (NTPV) (30). Genome sequencing and phylogenetic analyses revealed NTPV as a member of the Karimabad species complex with its closest genetic relative being Gabek Forest virus (GFV). Experiments showing productive replication in ruminant, rodent, sandfly, and human cell lines suggested a wide species tropism of NTPV (30). Additionally, seroprevalence studies at the site of sandfly collection as well as in other regions of Kenya confirmed the presence of neutralizing antibodies in humans and the potential of NTPV to infect humans (13.9% seropositivity, no cross-reaction with GFV) (30). Follow-up screenings of sandflies (9, 31) and also ticks (18) again detected NTPV, suggesting its continuous circulation in Kenya. However, no acute human infection has been described to date, so it remains unknown whether NTPV causes disease.

GFV was first isolated in 1961 from tissue pools of rodents, primates and other wild animals (32, 33). It is known to infect sandflies (34), and a serosurvey indicated human infections (29). GFV can produce a fulminating fatal illness in hamsters (35, 36) but its pathogenicity for humans is unknown since there are only serological data available.

The IFN system is the first line of defence of mammals against viruses. It is activated by pattern recognition receptors (PRRs) that are able to sense molecular structures (often nucleic acids) of the intruding pathogen. For example, the RNA “panhandle” formed by the 5’ and 3’ genome ends of incoming RVFV nucleocapsids is recognized by the cytoplasmic PRR Retinoic Acid-Inducible Gene I (RIG-I) (37) which subsequently interacts with the adapter protein Mitochondrial Antiviral Signaling (MAVS) via Caspase recruitment domains (CARDs) that are present on both RIG-I and MAVS (38). Activated MAVS then transmits the signal to the TANK-binding kinase 1 (TBK1) to phosphorylate the transcription factor IRF3 and trigger its import into the nucleus. IRF3 can transactivate the promoters of several IFN genes, especially of type I (IFN-α/β) and type III (IFN-λ), as well as other antiviral cytokines (39). The secreted IFNs bind to their cognate receptors on neighbouring cells where they set off the so-called JAK/STAT signalling cascade leading to the expression of IFN-stimulated genes (ISGs) (40). As many ISGs encode proteins with antiviral activity, phleboviruses have evolved the NSs proteins that counteract the induction of IFNs or disarm ISG activities (5–7). The quality and strength of such IFN evasion capabilities (“innate immunity phenotype”) can be helpful to assess the pathogenic potential of a virus.

Here, we provide first insights into the innate immunity phenotype of the phleboviruses NTPV and GFV. We show that both viruses can productively replicate in IFN-competent human cell lines that are derived from lung, liver and kidney material, and upregulate the transcription of IFNs, cytokines and ISGs to an extent that was lower than an IFN-inducing control virus lacking a functional NSs gene. We also establish that NTPV and GFV are sensitive to exogenously added IFNs, and that both these novel phleboviruses express NSs proteins that act as antagonists of the human IFN system by binding to the CARD of MAVS, thus inhibiting IFN induction. NTPV and GFV thus have the capacity to counteract the human IFN system.

## Results

To place our analyses in a context, we compared NTPV and GFV with two strains of the well-characterized phlebovirus RVFV, namely MP-12 (41) and Clone 13 (Cl13) (42). MP12 is an attenuated RVFV strain with a cell culture phenotype like virulent RVFV, since it expresses a functional IFN antagonist NSs. RVFV Cl13, by contrast, carries a large deletion in the NSs gene that turns it into a potent IFN inducer virus and lacks the ability to combat the antiviral host response (42, 43).

### Human cell line susceptibility to NTPV

The previous study by Tchouassi *et al.* on NTPV reported cell permissiveness and virus yields of cell lines from different mammalian species, and used the embryonic kidney line HEK293T as representative for humans (30). We expanded the human focus of such analyses with cell lines derived from human liver (Huh7), lung (A549 and H1299), and intestine (Caco-2), and compared them to the kidney line HEK293. We observed that NTPV established productive infection in all tested cell lines at 24 and 48 h post infection, both at an intermediate (0.1) (Fig. 1A and B) and low (0.01) (Fig. 2C and D) multiplicity of infection (MOI). High titers (approximately 10E6 or more plaque forming units [PFU]/ml at 48 h post infection (p.i.)) were reached by NTPV in Huh7, A549, H1299, and HEK293 cells, whereas titers in Caco-2 cells were lower by several orders of magnitude. GFV reached virus titers that tended to be slightly lower but were generally in the same range as those of NTPV in all cell lines except for Huh7 where it was reduced by about 1 log10 at the low MOI, and in Caco-2 where GFV remained in most cases undetectable. Both RVFV control strains, as expected, productively infected Huh7, A549, H1299, and HEK293 cell lines. In Caco-2 cells, reliable infection was established for both RVFV strains (as for NTPV) although at lower levels than in the other cell lines, but not for GFV. With this, we show a broad tropism of NTPV and GFV in cell lines from different human tissues, comparable to RVFV.

**Figure 1:**
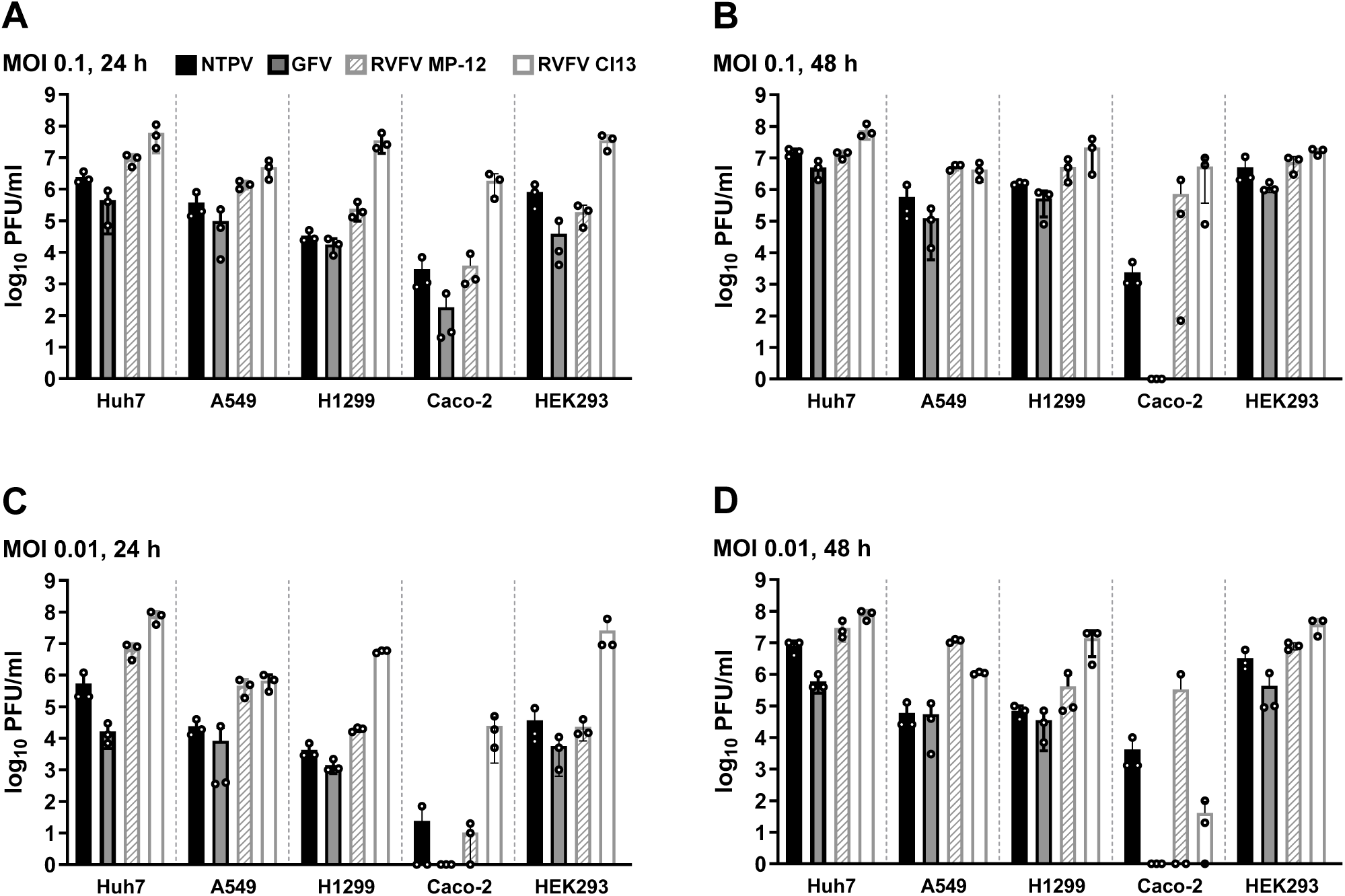
Human cell line susceptibility. Cells of the indicated lines were infected with NTPV, GFV, RVFV MP-12 or RVFV Cl13 at an MOI of 0.1 (A, B) and 0.01 (C, D) and virus titers were measured by plaque assay of supernatants at 24 h (A, C) and 48 h (B, D) p.i.. Individual titers (dots), geometric mean values (bars) and standard deviations from three technical replicates are shown.

**Figure 2:**
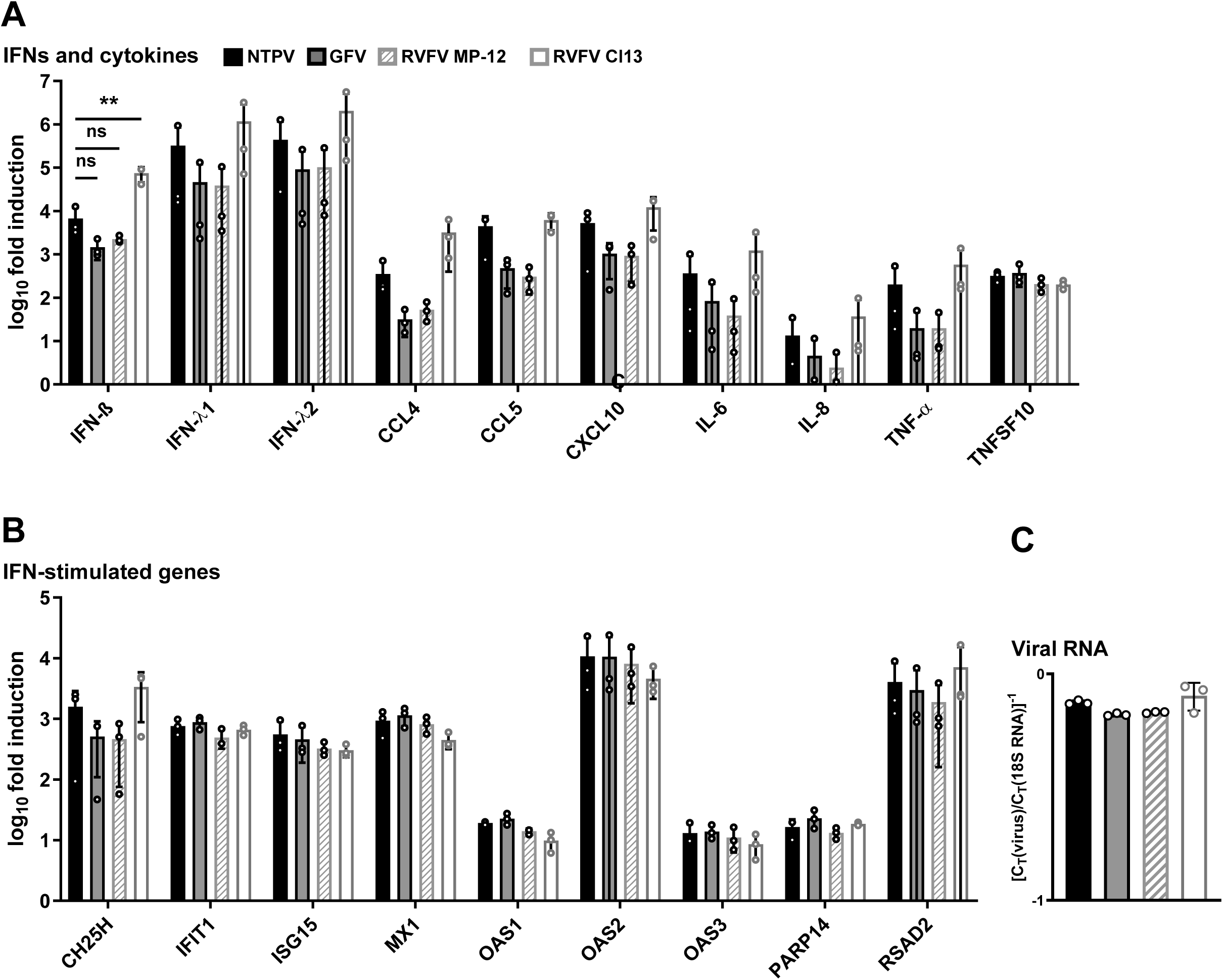
Expression of innate immune genes. Human A549 cells were infected with the indicated viruses at an MOI of 1 and harvested at 16 h post infection. Gene expression of select IFN and cytokine genes (A) and ISGs (B) was measured by quantitative real-time PCR analysis. Differential gene expression was calculated using the ΔΔCT method and results are given as fold-induction over the uninfected mock control. 18S rRNA was used as a reference gene. Viral gene expression (C) is pictured as CT value for viral L segment normalized to 18S rRNA control. Individual (dots), geometric mean values (bars) and standard deviations from three biological replicates are shown. Values were analyzed by one-way ANOVA with Dunnett’s post-test compared to NTPV. *P<0.0332; **P<0.0021; ns, not significant (Graphpad Prism). No label is added to the genes where statistical significance compared to NTPV was not reached.

### Innate immunity gene activation by phlebovirus infection

For efficient containment of virus infections, a rapid induction of IFNs and other cytokines and innate immunity genes is essential for the host, whereas their blockade benefits the virus (44, 45). Using the IFN-competent cell line A549 which was found to be permissive for all tested phleboviruses, we sought to examine the differential regulation of IFNs, cytokine genes, and ISGs by NTPV and GFV in relation to the two RVFV strains. Based on previous studies (46–52), we compiled a set of host genes indicative of virus infection. This set of marker genes includes representative interferons IFN-β, IFN-λ1 and IFN-λ2 and cytokines CCL4, CCL5, CXCL10 (IP-10), IL-6, IL-8 (CXCL8), TNF-α and TNFSF10 (TRAIL), as well as ISGs CH25H, IFIT1 (ISG56), ISG15, MX1, OAS proteins, PARP14 and RSAD2 (Viperin).

The human A549 cells were infected with an MOI of 1 for 16 h and subsequently analysed by RT-qPCR. Overall, NTPV infection elicited an innate immune response that was lower than the IFN-induction control RVFV Cl13, but tended to be slightly higher than in response to GFV and RVFV MP-12. All tested IFN and cytokine genes were most strongly induced by the NSs-deficient RVFV Cl13 as expected, followed by NTPV, and lowest for GFV and MP- 12, except for TNFSF10 which was induced to the same degree by all four phleboviruses (Fig. 2A). In contrast, ISGs were more equally induced with the exception of CH25H, where expression again tended to be higher upon NTPV and RVFV Cl13 infection (Fig. 2B). For wt phleboviruses (i.e. all bar Cl13), there was a trend towards higher innate immunity gene induction for NTPV compared to GFV or MP-12, but statistical significance was not reached. Notably, viral RNA loads (Fig. 2C) almost mirrored the gene induction patterns observed for the IFNs and cytokines except TNFSF10 (see Figure 2A), but not for most of the ISGs (see Figure 2B). Altogether, we show that NTPV induces the cytokines to levels that tend to be higher than those of GFV and RVFV MP12, but lower or similar than RVFV Cl13. Upregulation of ISGs, by contrast, was roughly equivalent to the other phleboviruses.

### Sensitivity to type I and type III IFN

IFNs signal through their respective receptors to induce a vast number of antiviral ISGs. We investigated the sensitivity of NTPV and GFV to type I and type III IFNs, which stimulate a similar subset of genes but differ in their receptor distribution: While type I IFNs (IFN-α/β) act on the IFN-α receptor (IFNAR) that is ubiquitously present on most cell types, type III IFNs (IFN-λ1 to 4) signal through the IFN-λ receptor (IFNLR) expressed almost exclusively on epithelial surfaces (39, 53). Compared to type I IFN, type III IFN induces a weaker but longer-lasting immune response. We pre-treated A549 cells for 16 h with IFN-α or IFN-λ and evaluated virus yields after 24 and 48 h of infection. Both NTPV and GFV, as well as the two RVFV strains that were used for comparison, were reduced by IFN-α pre-treatment, and at both 24 h and 48 h p.i. (Fig. 3A, B). Also IFN λ caused significant titer reductions of all tested viruses at 24 h p.i. (Fig. 3C). At 48 h p.i., RVFV MP-12 was able to overcome type III IFN action, while NTPV, GFV, and RVFV Cl13 growth was still significantly inhibited (Fig. 3D). These data show that NTPV and GFV are sensitive to exogenously added type I and III IFNs. Next, we examined whether the IFNs that are intrinsically induced by NTPV and GFV (see Fig. 2A) represent an obstacle for their replication. To do this, we used the FDA-approved drug Ruxolitinib, which interferes with type I and III IFN signalling by targeting IFN-receptor associated Janus kinases (JAK)1/2 (54). A549 cells were pre-treated with Ruxolitinib for 16 h, infected, and virus titers determined at 24 and 48 h p.i.. NTPV replication was boosted by Ruxolitinib at 48 h p.i. (Fig. 3E), comparable to the NSs-deleted control virus RVFV Cl13 (Fig. 3H). On the other hand, neither GFV (Fig. 3F) nor RVFV MP 12 titers (Fig. 3G) were elevated by Ruxolitinib in a statistically significant manner. However, the differences in Ruxolitinib dependency between the viruses appear small. Nonetheless, the data are in line with the tendency of NTPV to induce higher levels of IFNs than GFV (see Fig. 2), as they show that only NTPV benefits from blocking the action of endogenous IFN in a manner that is comparable to the IFN inducing control RVFV Cl13.

**Figure 3:**
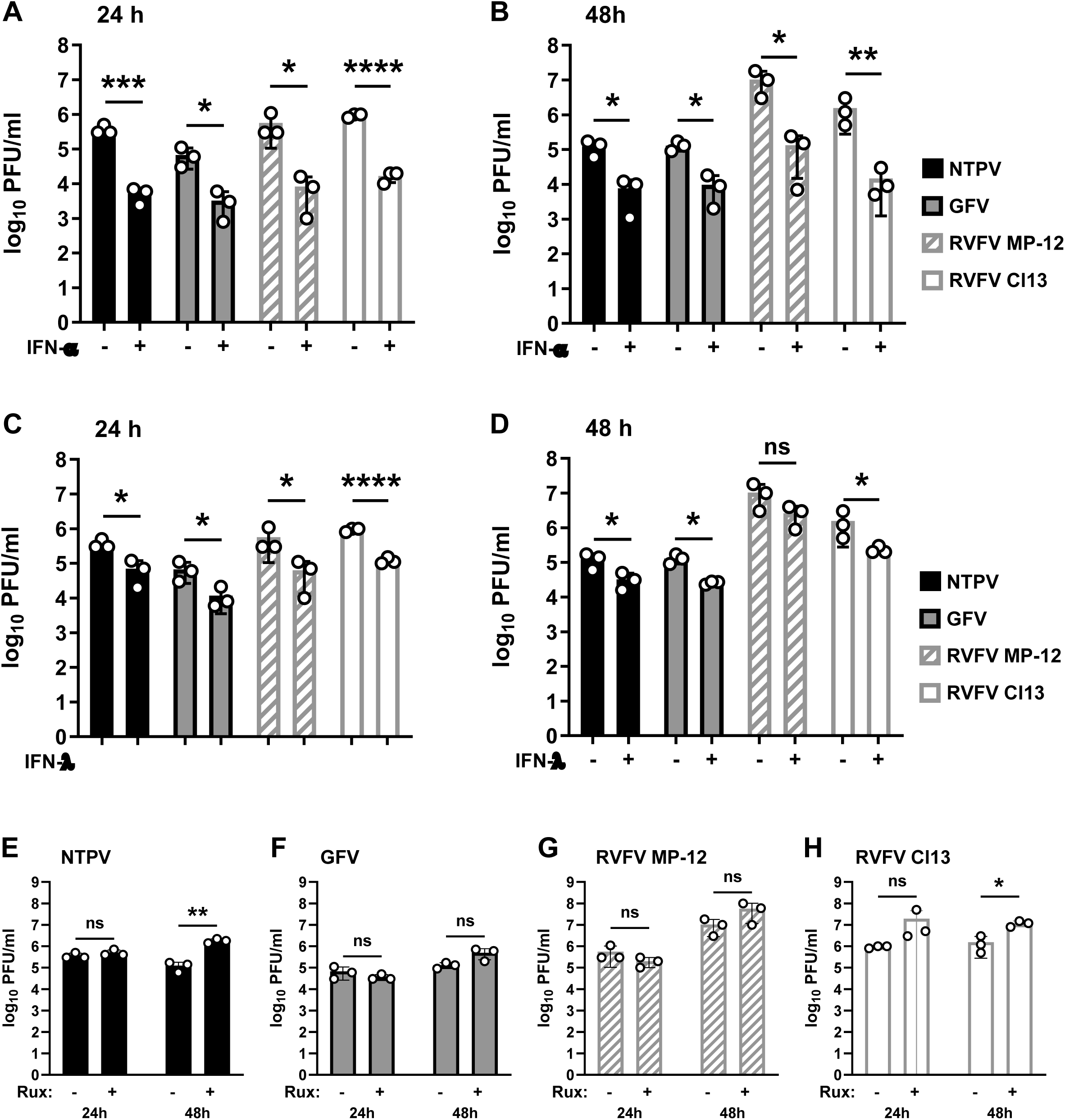
Sensitivity to antiviral interferons. A to D, IFN pretreatment. A549 cells were incubated with 1,000 U/ml IFN α(B/D) (A, B) or 100 ng/ml IFN λ3 (C, D) for 16 h and infected with the indicated viruses at an MOI of 0.01. Virus titers at 24 h (A, C) and 48 h (B, D) post infection were measured by plaque assay of supernatants. E to H, inhibition of IFN signaling. A549 cells were pre-treated with 1 μM Ruxolitinib (Rux) for 16 h, infected with the indicated viruses at an MOI of 0.01, and titers were determined at 24 and 48 h post infection by plaque assay titration of supernatants. Log-transformed titers were analyzed by unpaired one-tailed Student’s t test. *P<0.0332; **P<0.0021; ***P<0.0002; ****P<0.0001; ns, not significant.

### Inhibition of IFN induction and signalling by NTPV and GFV NSs proteins

For several other phleboviruses the non-structural protein NSs was identified as the virulence factor responsible for their anti-IFN potency (reviewed in (5–7)). To explore this for NTPV and GFV, we cloned and expressed their NSs genes and compared them to the well-described and highly potent RVFV NSs, which acts mainly by causing a general block of host mRNA transcription (42, 55). Moreover, since RVFV NSs might be an outlier in employing mostly proteasomal degradation and host cell transcription shutoff for its IFN-antagonistic function (56, 57), we also compared to the NSs of the phlebovirus SFSV, which acts primarily via stoichiometric inactivation of key antiviral host factors (58, 59). As negative controls, we included a plasmid coding for a C-terminally truncated, non-functional MxA protein, termed ΔMx (CTRL). In a first set of experiments, we investigated the effect of the NTPV and GFV NSs proteins on IFN induction, using a reporter construct with the firefly luciferase gene under control of the IFN-β promoter. Cells were co-transfected with the IFN-β promoter reporter and either of the expression constructs, stimulated using genomic RNA of Vesicular stomatitis virus (VSV) 24 h later, and then assayed for reporter gene activity after an additional 18 h incubation. Figure 4A shows that both NTPV and GFV NSs are able to counteract VSV RNA-stimulated IFN-β promoter activation, although not as strongly as RVFV NSs and SFSV NSs.

**Figure 4:**
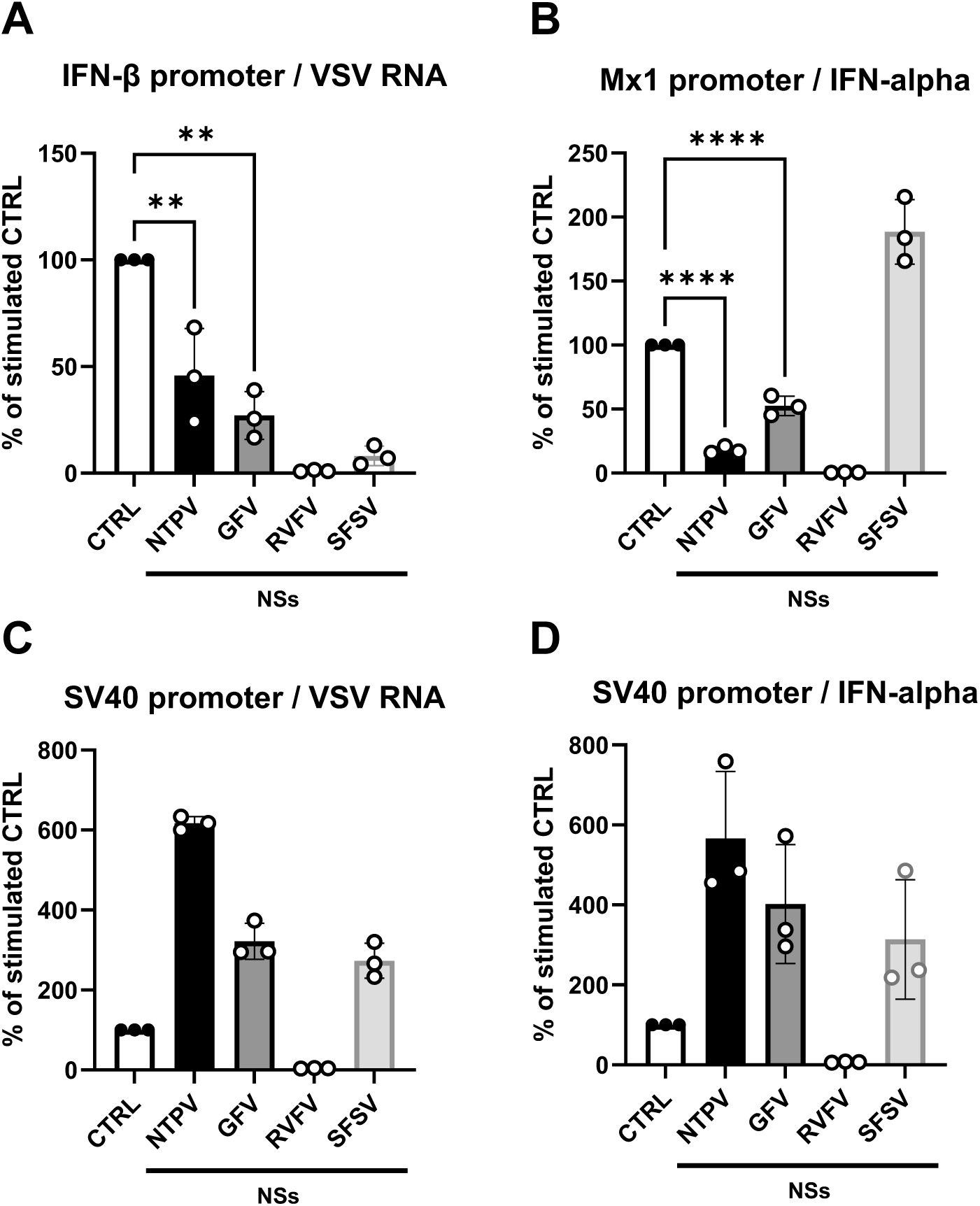
IFN-antagonism by NSs proteins. Human HEK293 cells were transfected with expression plasmids for the inactive control ΔMx (CTRL), untagged NSs of NTPV or GFV, or 3×FLAG-tagged NSs of RVFV or SFSV, as well as stimulation-dependent firefly luciferase and constitutively active *Renilla* luciferase reporters. At 24 h post transfection, promoter activation was induced by VSV-RNA transfection (50 ng/well) or IFN-α(B/D) (50 U/well). Cell lysates were harvested 18 h after stimulation for luciferase assays. Stimulation-dependent firefly luciferase reporters were derived from genes for IFN-β (A), or Mx1 (B), and the corresponding co-transfected constitutive SV40 promoter (C, D). Firefly and Renilla luciferase values of stimulated control samples were set to 100% within each biological replicate. Individual values (dots), geometric mean values (bars) and standard deviations from three biological replicates are shown. Values were analyzed by one-way ANOVA with Dunnett’s post-test compared to stimulated control, adjusted p values are shown for the IFN antagonism of NTPV and GFV NSs.

In a second set of experiments, we examined whether NTPV and GFV NSs could counteract IFN signaling, using a firefly luciferase reporter for the Mx1 promoter which exclusively responds to IFN (60). Cells co-transfected with the reporter and expression plasmids were stimulated with IFN-α for 18 h before reporter activity was measured. As seen in figure 4B, the NSs of both NTPV and GFV could significantly reduce IFN-dependent Mx1 promoter activation, but again the degree of inhibition was lower than for RVFV NSs. SFSV NSs did not antagonize Mx1 promoter activity, in line with our previous finding that it is unable to interfere with IFN signaling (61).

As mentioned, the NSs of RVFV inhibits the IFN system by a broad host transcription shutoff, whereas SFSV NSs sequesters specific host factors that are involved in antiviral defence. In order to clarify to which of these categories the NTPV and GFV NSs proteins belong to, our transfection mixes to measure IFN induction and IFN signaling (see Figs. 4A and B) also contained a Renilla luciferase reporter construct for the constitutively active SV40 promoter. As expected, the NSs of RVFV massively suppressed the SV40 promoter reporter, whereas the NSs of SFSV did not (Fig. 4 C and D). Moreover, both NPTV and GFV NSs behaved like SFSV, indicating the absence of a general host transcription shutoff. Taken together, our NSs overexpression/reporter assays indicate that NTPV and GFV NSs are capable of inhibiting IFN induction and, somewhat surprising for phleboviruses (5–7), also IFN signalling. Their inhibitory activities against innate immunity-related promoters are specific and not due to a general host cell transcription shutoff.

### NTPV and GFV NSs inhibit IFN induction at the level of MAVS

As far as it is known, inhibition of IFN induction is the most conserved activity of phleboviral NSs proteins (see data and references above). Whereas RVFV NSs blocks it at the level of mRNA synthesis by RNAP II (55–57), others target specific relays in the RIG-I-dependent antiviral signaling chain, e.g. SFSV preventing IFN promoter binding by IRF3 (58) or TOSV interacting with and degrading RIG-I (62).

In order to clarify the point of interference by NTPV and GFV NSs, we performed our IFN-β promoter assays by overexpression of either a constitutively active RIG-I mutant (RIG-I CARDs), MAVS, TBK1, or a constitutively active IRF3 mutant (IRF3 CA) to stimulate the RIG-I-dependent antiviral signaling chain at the different relay points. The data shown in figures 5A to D reveal that NTPV NSs was able to inhibit IFN-β promoter stimulation at the stages of RIG-I and MAVS, but not further downstream. GFV behaved similarly, albeit inhibition of MAVS signaling did not reach statistical significance. The NSs proteins of RVFV and SFSV, by contrast, inhibited all stages of the antiviral signaling chain, in agreement with their known mechanisms acting at the promotor level.

**Figure 5:**
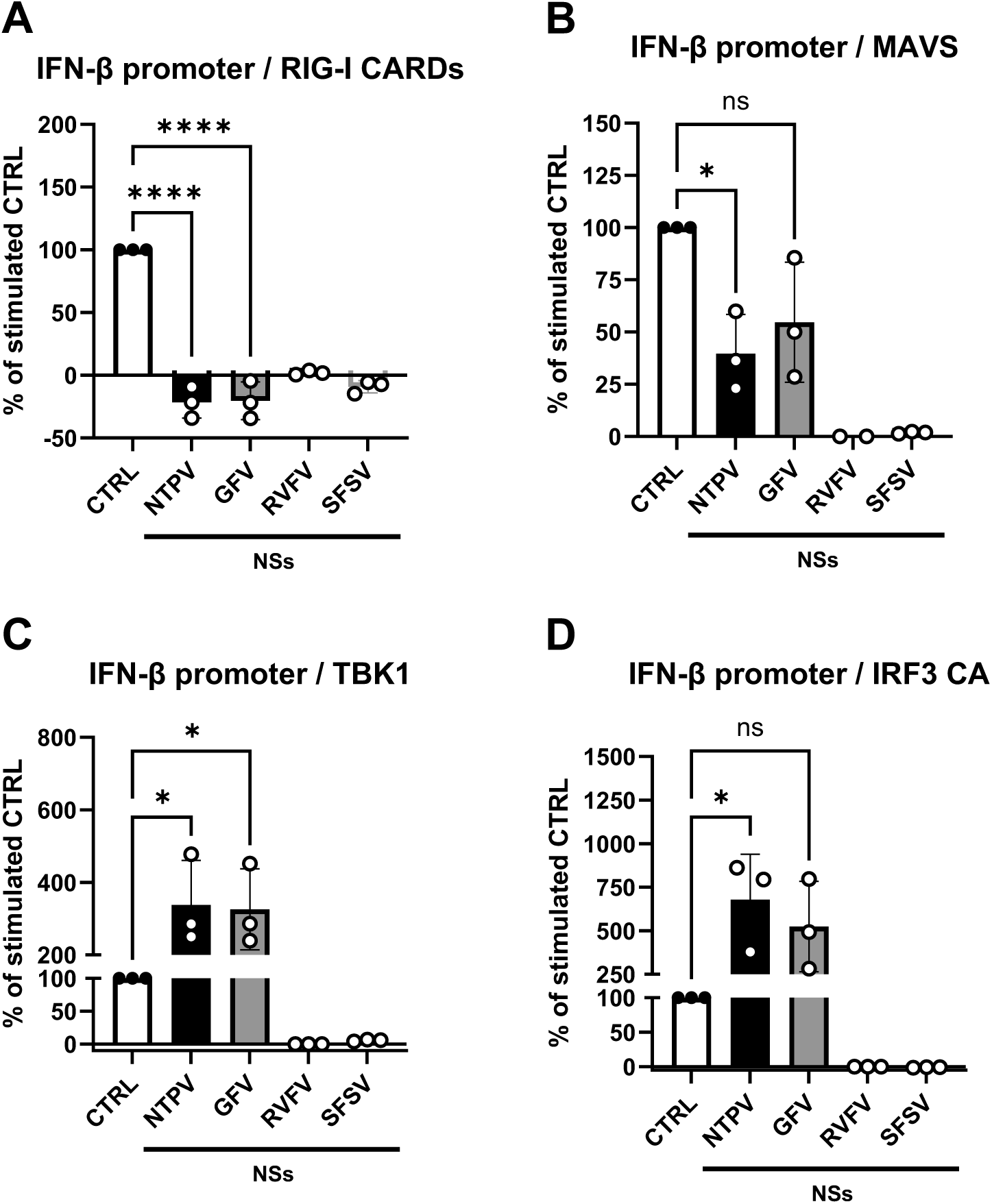
Identification of the antiviral signaling step targeted by NSs proteins. HEK293 cells were transfected with IFN-β reporter and expression plasmids as described for figure 4, and additionally transfected with expression plasmids either for human RIG-I CARD (A), MAVS (B), TBK1 (C), or the constitutively active IRF3(5D) mutant (IRF3 CA; D). Statistical analysis was performed as described for figure 4.

### NTPV and GFV NSs directly interact with the MAVS CARD

In order to investigate whether the observed MAVS inhibition is due to protein-protein interaction, we employed the NanoBiT complementation system in which a NanoLuc reporter protein is artificially split into two subunits (18 kDa LgBiT ánd 1.3 kDa SmBiT) that only assemble to a functional luciferase when both are fused to proteins that interact with each other (63) (Fig. 6A). Unlike the traditional co-immunoprecipitation assays, the NanoBiT system measures protein-protein interactions within the cellular environment, is easily quantifiable, and ignores interactions that are not direct. We constructed cDNA plasmids in which the NSs proteins of SFSV, NTPV, and GFV are fused to the nanoLuc LgBiT, and human IRF3 and MAVS to the SmBiT. SFSV NSs and IRF3 thereby served as specificity controls and their interaction (58) as additional positive control. Human HEK293T cells were transfected with different combinations of those cDNA in parallel to the positive and negative control plasmids provided with the NanoBiT system. As seen in figure 6B, SFSV NSs expectedly produced a NanoLuc complementation signal when co-expressed with the IRF3 construct, whereas co-expression with the provided negative control (HaloTag) or with MAVS did not. Contrary to that, both NTPV and GFV NSs produced only a signal with MAVS, but not with IRF3. These data confirm our previous conclusions on the SFSV NSs- IRF3 interaction (58) by an independent method, and demonstrate a direct and specific interaction of NTPV and GFV NSs with MAVS.

**Figure 6:**
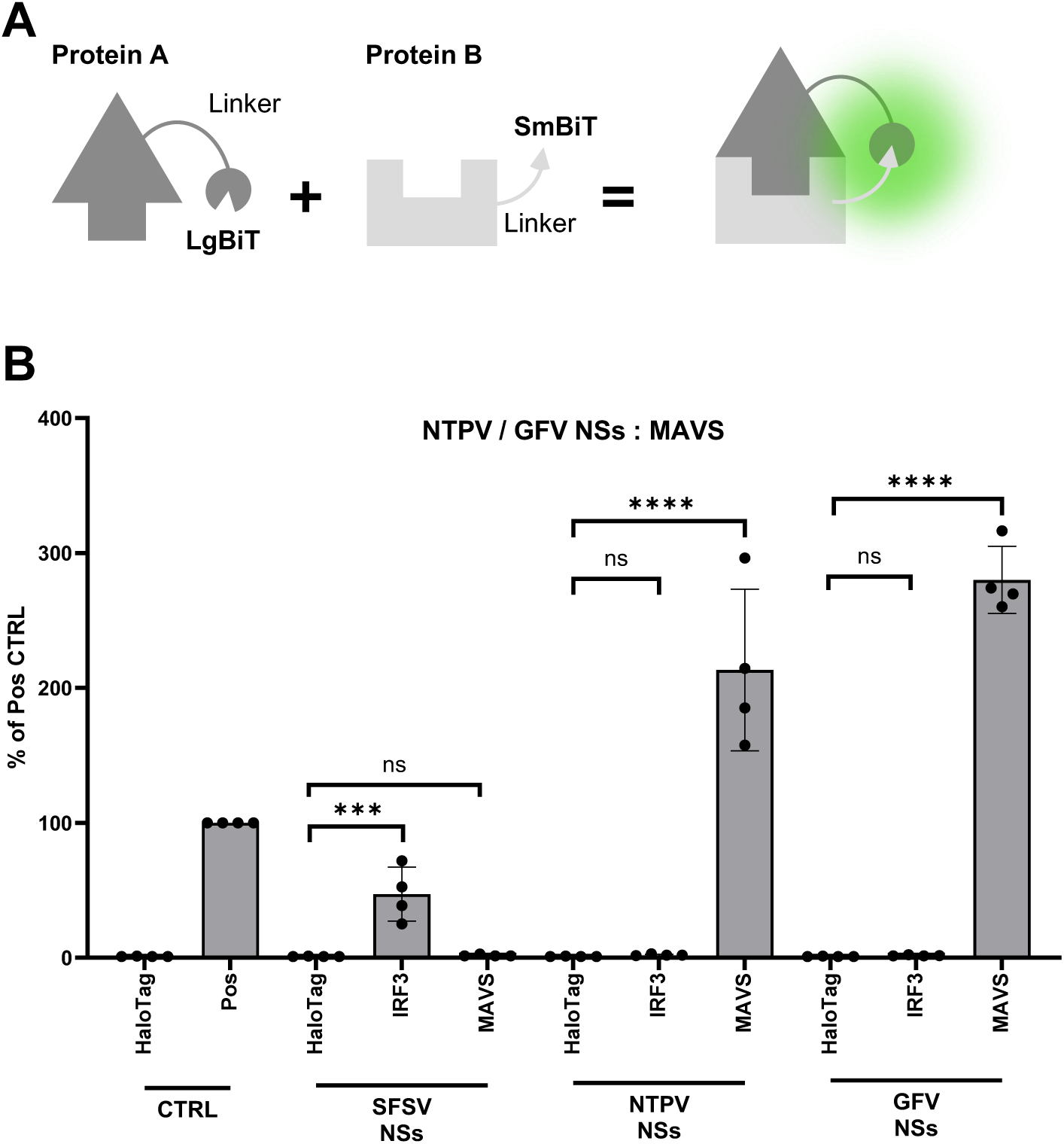
Testing direct interaction of NSs proteins with MAVS in the NanoBiT system. (A) Cartoon showing the principle of the NanoBiT complementation system. (B) NanoBiT interaction assay. HEK293T cells were transfected with expression constructs pBiT1.1- N[TK/LgBiT]-SFSV-NSs, pBiT1.1-N[TK/LgBiT]-NTPV-NSs, or pBiT1.1-N[TK/LgBiT]- GFV-NSs encoding the respective NSs proteins fused to the nanoLuc LgBiT combined with either pBit2.1-N[TK/SmBit]-MAVS or pBit2.1-N[TK/SmBit]-IRF3 encoding cellular proteins fused to the SmBit or with a HaloTag fused to SmBiT as negative control. Positive control plasmids LgBiT-PRKAR2A and SmBiT-PRKACA were transfected in parallel (Pos). Statistical analysis was performed as described for figure 4.

We further exploited the NanoBiT system to map the site on MAVS targeted by NTPV and GFV NSs. The 540 amino acids-long MAVS consists of an N-terminal CARD (responsible for interaction with RIG-I), followed by a proline-rich domain (PRR) and a long linker region (mediating downstream signaling), and a C-terminal transmembrane (TM) domain (Fig. 7A, top) (64, 65). We generated a series of C-terminal, N-terminal and internal MAVS deletion mutants fused to the SmBiT to test them in the NanoBiT system for NSs interactions (Fig. 7B). The mutants lacking the N terminus (miniMAVS, ΔN101) lost the capacity to interact with NTPV and GFV NSs, whereas mutants retaining the N terminus were still interacting. Even when just the N-terminal CARD was used there was a signal above background, which however did not reach significance level in the multiple comparisons. Although the level of interaction of the CARD and some of the other mutants containing the CARD was lower than for full-length MAVS, these data clearly show that the interactions of NTPV and GFV NSs are mediated by the CARD of MAVS.

**Figure 7:**
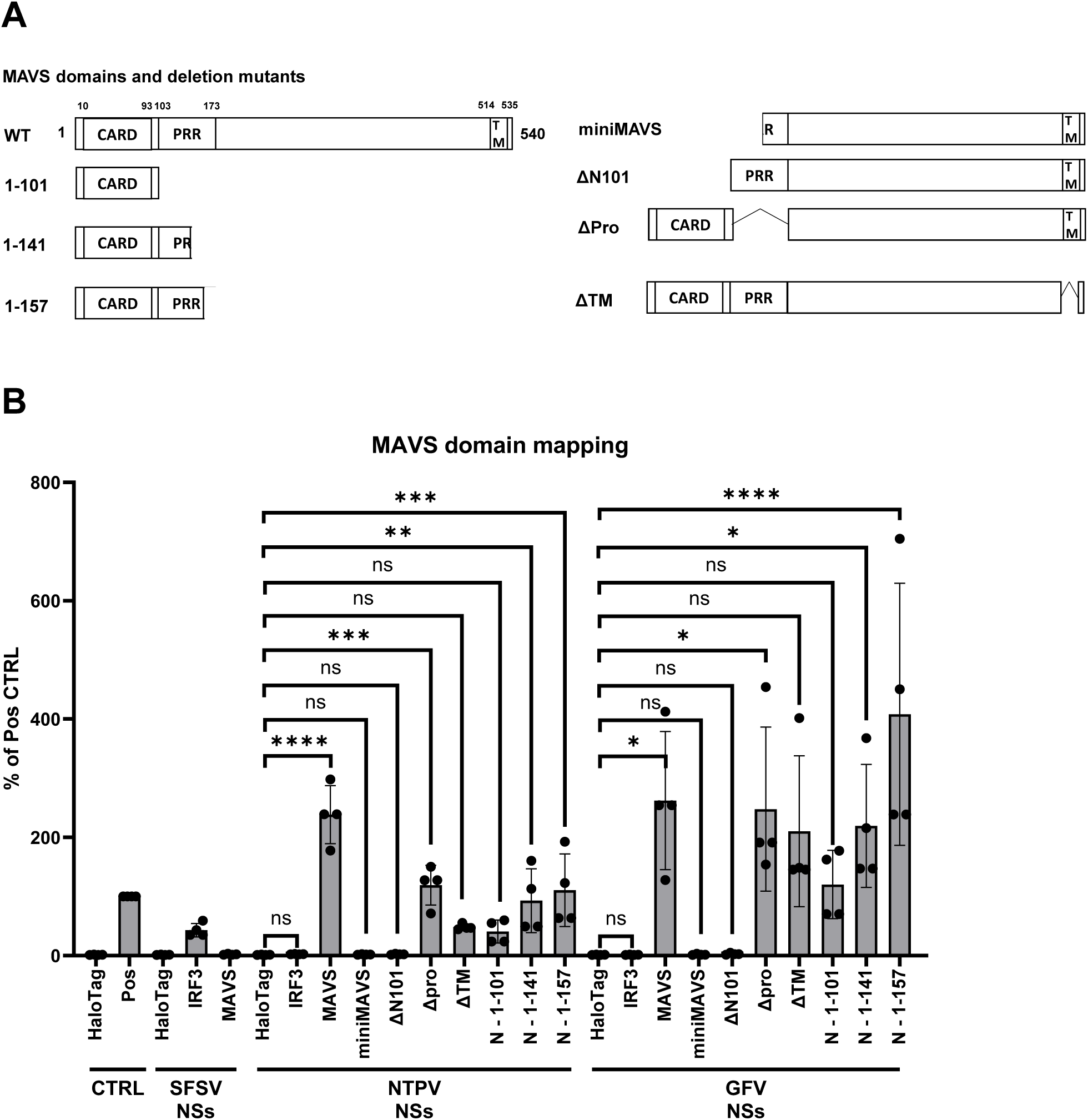
Mapping the NSs interaction domain on MAVS. (A) Schematic representation of the MAVS domains and the tested deletion mutants. (B) NanoBiT interaction assay. HEK293T cells were transfected with either of the LgBiT-fused NSs plasmids from figure 6 in combination with one of the plasmids expressing SmBiT-fused deletion mutants of MAVS (see A). Statistical analysis was performed as described for figure 4.

## Discussion

The ever-growing genus *Phlebovirus* (family *Phenuiviridae*, class *Bunyaviricetes*) currently comprises 82 named species (4, 66), but for most members the pathogenic potential is unknown. Viral zoonotic risk prediction can be divided into the assessment of (i) the capacity to infect humans (spillover) and (ii) the virulence once the virus has entered the human host (67–69). A risk assessment study employing machine-learning models predicted a high zoonotic potential for several phleboviruses including NTPV and GFV (70). In line with this, NTPV has some seroprevalence in the Kenyan population (30), and the related GFV (formerly known as SudAn 754-61) is seroprevalent in Sudan, Egypt and Nigeria (29). For both viruses we observed a high permissiveness of human cell lines derived from liver, lung, and kidney, but a low permissiveness of an intestinal line. Thus, NTPV and GFV may have the potential to cause viremia and infect inner organs. In any case, both viruses exhibited cell culture features compatible with a human pathogenic virus. The degree of virulence cannot be assessed from these data, but given the absence of noted disease it may not be high. It needs however be remarked that the IFN sensitivity of NTPV and GFV is in the range of 2 log10 steps reduction after treatment with 1,000 U/ml IFN-α, which is similar to RVFV but lower than what we observed for the successful human viruses SARS-CoV-1, SARS-CoV-2, and MERS-CoV (71, 72).

Viral antagonism of IFN induction and antiviral IFN action by a particular virus is an important parameter of risk assessment, as has recently been exemplified by SARS-CoV-2 variants of concern (71, 73, 74). Regarding such an innate immunity profile, we observed some slight differences between the closely related NTPV and GFV. NTPV tended to induce higher mRNA levels of IFNs and other cytokines than GFV, but these were still lower compared to the IFN-inducing control virus RVFV Cl13 lacking a full-length NSs. In general, IFN induction by GFV was closer to RVFV MP-12, whereas for NTPV it was in between the IFN suppressor MP-12 and the IFN inducer Cl13. IFN induction or its suppression plays a vital role for health vs. disease in the infected. The IFN inducing Cl13 mutant of RVFV is highly attenuated in wt mice, but can kill animals with a deficient IFN system within 40 hours, whereas the virulent strain ZH548 also kills IFN-competent wt mice (75). Similarly, the low-pathogenic strain Balliet of the phlebovirus Punta Toro virus (PTV) – but not the more virulent strain Adames – is an inducer of IFNs, cytokines and ISGs, and the high cytokine levels were associated with a protective effect against disease (76, 77). Moreover, the related Uukuniemi virus (UUKV; genus *Uukuvirus*, family *Phenuiviridae*) is believed to be apathogenic for humans (78), and again has very limited capacity to down modulate IFN induction (79). Thus, for members of the *Phenuiviridae* there are differences in IFN and cytokine upregulation that can have an impact on virulence.

In the case of NTPV and GFV, both viruses were able to inhibit the human IFN system. Their NSs proteins acted on the induction of IFN as well as on the signaling by exogeneous IFN, albeit there were minor differences. The slightly weaker inhibition of IFN induction by the NSs of NTPV seems in line with the higher IFN induction by NTPV and the observation that NTPV benefited from inhibition of the virus-induced IFN signalling by Ruxolitinib whereas GFV growth was not enhanced by the pharmacological blockade of IFN signaling. However, the differences between the viruses are relatively minor. Interestingly, there were clear differences when compared to the other phleboviruses RVFV and SFSV. Firstly, NTPV and GFV did not impose a host cell shutoff like RVFV NSs did. This indicates they target specific components of the IFN system, unlike RVFV which targets the host transcription machinery (7). In that respect they were similar to SFSV and many other phlebo- and phenuiviruses (5–7). Secondly, the NSs of NTPV and GFV were active against both IFN induction and IFN signaling, which distinguishes them from SFSV which inhibits IFN induction, but not IFN signaling (61). Thus, NTPV and GFV resemble the relatively benign SFSV in that their NSs is an IFN-specific inhibitor, but like the more virulent RVFV they interfere with both phases of the IFN response, albeit not as strongly. From this innate immunity profile one can conclude that the virulence of the two novel and related phleboviruses is not as high as of RVFV.

IFN induction inhibition seems to be the most conserved activity of phlebovirus NSs proteins (see above and (5–7)). Therefore, to gain further information for the two novel phleboviruses, we investigated the affected infection step and identified MAVS as the relevant host cell interactor. MAVS is the key signaling adapter for intracellular RNA virus sensing (64, 65), and therefore targeted by many viral antagonists (80, 81). Some viruses degrade it such as hepatitis C virus which cleaves MAVS via its NS3/4a protease (82) or SARS-CoV-2 which induces its proteasomal digestion by its NSP5 protein (83). Other viral proteins interact stoichiometrically with MAVS to block downstream signaling, e.g. influenza virus PB2 (84), Dengue virus (DENV) NSP4 (85) or Zika virus (ZIKV) NS4A (86). With regards to phenuiviruses, a (comparatively weak) MAVS interaction with NSs has only reported for UUKV (genus Uukuvirus) (79), whereas, to our knowledge, not for any member of the genus phlebovirus. Also non-phleboviral phenuiviruses tend to not target MAVS, but rather TBK1 and other factors of IFN induction (5–7).

We found that NPTV and GFV NSs specifically and strongly bind to the CARD of MAVS. Such an occupation of the MAVS CARD is expected to result in a block of CARD-CARD interaction with activated RIG-I. Indeed, our reporter data indicate that both NSs proteins efficiently block IFN induction by the CARD of RIG-I as well as by MAVS overexpression (although less strongly), whereas IFN induction by TBK1 or IRF3 was unaffected. The fact that the NSs proteins cannot completely block IFN induction by overexpressed MAVS is most likely due to a partial stoichiometric outcompeting of NSs by the elevated levels of MAVS. So far, we only could identify DENV NSP4 and ZIKV NS4A (85, 86) as other viral IFN antagonists that bind the MAVS CARD (together with the TM domain in these cases) (85). It can however be expected that other viral MAVS interactors which were so far not characterized further would also bind the CARD. Our experiments were performed with human MAVS, and it should be noted that the natural reservoir hosts of NTPV and GFV are not known. The diversity of MAVS protein sequences from different animal species is relatively high, and the CARD is the most conserved domain (87), although some of its sites underwent positive selection (88). Thus, the CARD is not only an ideal target for viral antagonism due to its central role in signaling, but also because of its high sequence conservation. An IFN antagonism based on the MAVS CARD interaction may be one of the features that enable a broad host tropism.

To conclude, our data demonstrate that the novel phleboviruses NTPV and GFV, for which seroepidemiological data indicate human infections, can productively infect human cells from a range of inner organs, and handle the human innate immune system by suppressing IFN induction and IFN signaling through their NSs proteins. NTPV and GFV may thus have potential to further adapt to humans.

## Materials and Methods

### Cells and viruses

A549, BHK, H1299 (kindly provided by Marcel Müller, Charité Universitätsmedizin Berlin, Germany), HEK293, HEK293T, Huh7, VeroE6, and Vero76 cells were cultivated in cell culture medium CCM34 (Dulbecco’s modified Eagle’s medium (DMEM) with addition of 17.8 mg/l L-alanine, 0.7 g/l glycine, 75 mg/l L-glutamic acid, 25 mg/l L-proline, 0.1 mg/l biotin, 25 mg/l hypoxanthine, and 3.7 g/l sodium bicarbonate) supplemented with 10% heat-inactivated fetal calf serum (FCS; Thermo Fisher Scientific), and 5% Penicillin/Streptomycin/Glutamine (PSQ; Thermo Fisher Scientific) in a 5% CO_2_ atmosphere at 37°C. Caco-2 cells (kindly provided by Eric Miska, Gurdon Institute and Department of Genetics, University of Cambridge, UK) were cultivated in CCM34 supplemented with 15% FCS and 5% PSQ in a 5% CO_2_ atmosphere at 37°C. RVFV Clone 13 (Cl13) and NTPV strain MRG54-KE-2014 (30) were propagated on VeroE6 cells. RVFV strain MP-12 was propagated on BHK cells. GFV isolate Sud AN 754-61 was propagated on Vero76 cells. For all virus stocks, cell supernatant was collected 3 days post infection and centrifuged for 5 min at 800∈g to remove cellular debris. Virus stocks were titrated by plaque assay on BHK cells with 3 (RVFV) or 4 days (NTPV, GFV) incubation.

### Plasmids

To generate expression plasmids encoding untagged NSs of NTPV (GenBank accession no. MF695811.1) and GFV (KF297905.1), the viral gene sequences were amplified from cDNA and inserted into pI.18 via its 5ʹ *Bam*HI and 3ʹ *Xho*I restriction sites. Primer sequences are available on request. Plasmids pI.18-RVFV-NSs-3×FLAG and pI.18-3×FLAG- ΔMx were previously described (57, 58), and pRL-SV40 was purchased from Promega. Firefly luciferase reporter constructs p-125Luc (89)and pGL3-MX1P (90) were kindly donated by Takashi Fujita and Georg Kochs, respectively.

Expression plasmid pCDNA3.1-TOPO-RIG-I CARD was cloned by PCR-amplifying the coding region of human RIG-I from aa 1 to 284. Plasmids for human MAVS was kindly provided by Shizuo Akira (91) and plasmids for human TBK1 (92) and IRF3(5D) (93)were kindly provided by John Hiscott.

For construction of the NanoBit plasmids, the ORFs encoding the NSs proteins of SFSV, NTPV and GFV were cloned into the LgBit fusion vector, pBiT1.1-N[TK/LgBiT] (NanoBiT PPI MCS Starter System Vectors, Promega) between the *Eco*RI and *Nhe*I restriction sites using In-Fusion Snap Assembly (Takara) recombination according to the manufacturer’s protocol, resulting in N-terminally LgBiT-tagged pBiT1.1-N[TK/LgBiT]-SFSV-NSs, pBiT1.1-N[TK/LgBiT]-NTPV-NSs, and pBiT1.1-N[TK/LgBiT]-GFV-NSs. ORFs encoding MAVS and IRF3 were similarly cloned into the SmBit fusion vector pBiT2.1-N[TK/SmBiT] (Promega) between EcoRI and NheI sites using the same recombination method to produce N-terminally SmBiT-tagged pBit2.1-N[TK/SmBit]-MAVS and pBit2.1-N[TK/SmBit]-IRF3. Derivatives of pBit2.1-N[TK/SmBit]-MAVS were produced using the In-Fusion Snap Assembly recombination to remove the coding sequence corresponding to amino acids 1-141 (pBit2.1-N[TK/SmBit]-miniMAVS), 1-100 (pBit2.1-N[TK/SmBit]-MAVS-ΔN101), 103-173(pBit2.1-N[TK/SmBit]-MAVS-Δpro), 514-535 (pBit2.1-N[TK/SmBit]-MAVS-ΔTM), 102-540 (pBit2.1-N[TK/SmBit]-MAVS-N-1-101), 142-540 (pBit2.1-N[TK/SmBit]-MAVS-N1-141), 158-540 (pBit2.1-N[TK/SmBit]-MAVS-N1-157).

NanoBiT® Negative Control Vector encoding the HaloTag with a C-terminal SmBiT tag and the positive control plasmids LgBiT-PRKAR2A and SmBiT-PRKACA were purchased from Promega.

### Virus growth on human cell lines

Cells seeded into 24-well plates (5∈10^4^ per well) were infected the following day at a multiplicity of infection (MOI) of 0.1 or 0.01. At the indicated times post infection, cell supernatants were collected and titrated by plaque assay on BHK cells.

### Differential gene expression of innate immunity genes

Cells seeded in 12-well plates (1.5∈10^5^ per well) were infected the following day at an MOI of 1. At 16 h post infection, cells were lysed in 350 µl RLT buffer (Qiagen) supplemented with β-mercaptoethanol (1:100). RNA was extracted using the RNeasy mini kit (Qiagen) and reverse transcribed into cDNA using the prime Script RT reagent kit (Takara). Differential regulation of cellular genes was assessed using TB Green™ Premix Ex Taq™ II (Tli RNase H Plus; Takara) according to manufacturer’s instructions with QuantiTect primer assays (Qiagen; Hs_CCL4_1_SG, QT01008070; Hs_CCL5_1_SG, QT00090083; Hs_CH25H_1_SG, QT00202370; Hs_CXCL10_1_SG, QT01003065; Hs_CXCL8_1_SG, QT00000322; Hs_IFIT1_1_SG, QT00201012; Hs_IFNB1_1_SG, QT00203763; Hs_IFNL1_2_SG, QT01033564; Hs_IFNL2_1_SG, QT00222488; Hs_IL6_1_SG, QT00083720; Hs_ISG15_1_SG, QT00072814; Hs_MX1_1_SG, QT00090895; Hs_OAS1_1_SG, QT00099134; Hs_OAS2_1_SG, QT01005256; Hs_OAS3_1_SG, QT01005277; Hs_PARP14_1_SG, QT00087444; Hs_RR18s, QT00199367; Hs_RSAD2_1_SG, QT00005271; Hs_TNF_3_SG, QT01079561; Hs_TNFSF10_1_SG, QT00079212). Fold gene induction was calculated according to the threshold cycle (ΔΔC_T_) method using 18S rRNA as a reference gene (94). Viral load in samples was assessed using Premix Ex Taq (probe qPCR; Takara) according to manufacturer’s instructions with primers and probes as follows: for NTPV L, fwd, 5’- GCAAGAAAGCACTGTGGTGG-3’, rev, 5’-CGTATGATGATCGGCCACCA-3’, probe, 5’-6-FAM-ACAGCCACCTCTGATGATGC-BHQ1-3’ (30); for GFV L, fwd, 5’- GCAAGAAAACACTGTGGTGG-3’, rev, 5’-CGGATTATGATGGGCCACCA-3’, probe, 5’-6-FAM-ACAGCCACCTCGGACGATGC-BHQ1-3’ (designed to correspond to the NTPV primers); for RVFV L, fwd, 5’-TGAAAATTCCTGAGACACATGG-3’, rev, 5’- ACTTCCTTGCATCATCTGATG-3’, probe, 5’-6-FAM CAATGTAAGGGGCCTGTGTGGACTTGTG BHQ1-3’ (95).

### Inhibitor treatment

Cells seeded into 24-well plates (5∈10^4^ per well) were pre-treated for 16 h with 1000 U/ml pan-species IFN-α(B/D) (PBL Assay Science) (96), 100 ng/ml purified recombinant IFN-λ3 (kind gift from Rune Hartmann, Department for Molecular Biology and Genetics, Aarhus University, Aarhus, Denmark) (97), or with 1 μM Ruxolitinib (Selleckchem) and infected at an MOI of 0.01. At the indicated times post infection, cell supernatants were collected and titrated by plaque assay on BHK cells.

### Dual luciferase assay

HEK293 cells seeded into 96-well plates (1.5∈10^4^ per well) were transfected the following day with firefly and *Renilla* luciferase reporter constructs (40 ng each), as well as expression constructs for NSs proteins or the control protein ΔMx (10 ng) using TransIT-LT1 (Mirus Bio LLC). 24 h post transfection, cells were either treated with 50 U/well IFN-α(B/D) (PBL Assay Science) or transfected with 50 ng viral RNA (isolated from Vesicular Stomatitis virus (VSV) by phenol-chloroform extraction frLLCom PEG8000- precipitated VSV particles (98) using Endofectin (GeneCopoeia). Luciferase activities were measured 18 h later using a Dual-luciferase reporter assay system (Promega) according to the manufacturer’s instructions. Firefly luciferase values of unstimulated control samples were subtracted from values of stimulated samples, and the resulting values for the ΔMx control were set to 100% within each biological replicate.

### NanoBiT protein–protein interaction assay

HEK293T cells were seeded at 1×10⁴ cells per well into white clear bottom 96-well plates (Corning) in Opti-MEM medium (Thermo Fisher) supplemented with 100 U/ml penicillin and 100 µg/ml streptomycin (Gibco). After 24 hours, cells were transfected in duplicate, with 50 ng of one LgBiT fusion construct in combination with either a candidate interaction partner or the negative control plasmid encoding HaloTag fused to C-terminal SmBiT (NanoBiT® Negative Control Vector; Promega), using TransIT- LT1 transfection reagent (Mirus Bio LLC). The control plasmids LgBiT-PRKAR2A and SmBiT-PRKACA (Promega) were included in all assay as a positive control as a known strong interactor reference. After 24 h, reconstituted Nano-Glo Live Cell Reagent (Promega) was added directly to each well. Plates were incubated for 15 mins at 37 °C, after which luminescence was measured using a Spark multimode plate reader (Tecan).

## Acknowledgements

We are grateful to Takashi Fujita, Georg Kochs, Shizuo Akira, and John Hiscott for kindly providing plasmid constructs, and Anna Hoffbauer and Besim Berisha for technical assistance.

Work in the F.W. laboratory is funded by the Deutsche Forschungsgemeinschaft (DFG) SFB 1021 (project number 197785619) and GRK 2355 (project number 325443116), and the Swedish Research Council (project number 2018-05766).

## CRediT authorship contribution statement

MJP: conceptualization, data curation, formal analysis, investigation, methodology, validation, visualization, writing – review & editing

UF: conceptualization, data curation, formal analysis, investigation, methodology, validation, visualization, writing – original draft, writing – review & editing validation

DPT: resources, writing – review & editing

SJ: conceptualization, data curation, resources, writing – review & editing

FW: conceptualization, data curation, formal analysis, funding acquisition, project administration, supervision, visualization, writing – original draft, writing – review & editing

